# Ribosomes are covered by a coat of flexible tails of ribosomal proteins

**DOI:** 10.64898/2026.06.18.733196

**Authors:** Hugo McGrath, Rudolf Kvasňovský, Xingyue Guan, Michal Kolář

## Abstract

The three dimensional architecture of ribosomes is known to atomic detail thanks to X-ray crystallography or cryogenic electron microscopy. However, many ribosomal proteins contain terminal regions that are averaged out during electron density reconstructions, rendering them absent from experimental models. These flexible protein tails (FPTs) collectively form an invisible coat on the ribosome surface that has been systematically overlooked. Here we assessed 39 experimental models of ribosomes spanning bacteria, eukaryotes, and mitochondria, and analysed their FPTs using various bioinformatic tools. We show that mitoribosomes have the highest proportion of proteins carrying FPTs, as well as the longest and most numerous FPTs. Structural predictions indicated that FPTs are predominantly disordered. Comparison with proteome-wide background frequencies revealed class-specific compositional biases. Depletion of aromatic residues in all classes and sequence-based predictions argue against an intrinsic propensity of FPTs for protein-driven phase condensation. Compositional patterns point to roles for FPTs in RNA contacts. Our work provides a comprehensive overview of the understudied and likely functional part of the ribosomes.

## 1 Introduction

The ribosome is the universal molecular machine that decodes messenger RNA and synthesises proteins. Decades of X-ray crystallography and, more recently, cryogenic electron microscopy (cryo-EM) have yielded high-resolution structures of ribosomes from all domains of life, revealing in atomic detail the mechanisms of decoding, peptide bond formation, tRNA and nascent-chain translocation, and the binding of numerous translation factors. ^1,2^ Yet these structures share a systematic limitation: they capture only those parts of the ribosome that are sufficiently ordered to be resolved.

Comparing the sequences of ribosomal proteins (r-proteins) with their experimental structures, we found that substantial portions of many r-proteins are missing. These mostly terminal regions are likely disordered in crystals and averaged out during cryo-EM image processing, and therefore remain unresolved. We term them *flexible protein tails* (FPTs) and argue that, collectively, they form an invisible protein coat on the ribosome surface. Their extent, sequence properties and potential functions have not yet been systematically characterised.

Intrinsically disordered proteins (IDPs) and regions (IDRs) are now recognised as a functionally important and abundant component of proteomes across all domains of life. ^3–5^ Unlike folded globular domains, IDRs lack a stable three-dimensional structure under physiological conditions reflecting the rugged Gibbs energy hypersurface with many equienergetic minima. Still, they mediate a wide range of biological functions, including protein–protein and protein–nucleic acid recognition, post-translational modification, and the formation of membraneless organelles via liquid–liquid phase separation (LLPS) or more broadly biomolecular condensation.^6,7^ Their amino acid composition is characteristic and distinct from that of structured proteins: IDPs and IDRs are enriched in polar, charged and small residues, and depleted in bulky hydrophobics. This composition arguably encodes their biophysical properties and functional roles.^8^ Ribosomal proteins of the conserved core are known to carry extensions and insertions that are more prevalent in eukaryotes than in bacteria, ^9^ yet the properties of those experimentally unresolved regions remain elusive. While FPTs may share compositional features with IDRs and are often disordered by the criteria established for IDRs, they are defined here operationally by their absence from experimental structural models. This criterion encompasses genuinely disordered sequences as well as regions that may retain partial secondary or tertiary structure but remain flexibly tethered to the ribosome surface.

The vast majority of mitoribosomal proteins are nuclear-encoded, synthesised in the cytosol, and imported into the mitochondrion,^10^ and many carry a cleavable N-terminal mitochondrial targeting sequence (MTS) that is removed by the mitochondrial processing peptidase (MPP) upon import.^11^ These MTSs overlap with many of the FPTs and do not remain a part of the assembled mitoribosome.

Here we catalogue the FPTs on the surface of ribosomes spanning bacteria, eukaryotes, and mitochondria. We find that these regions differ in prevalence and length across ribosome types, with mitoribosomes having the highest proportion of proteins carrying FPTs as well as the longest and most numerous FPTs, and that FPTs are predominantly disordered in mitochondrial and eukaryotic ribosomes, whereas bacterial ribosomes are more heterogeneous. We further show that the FPT amino acid composition deviates from proteome-wide averages differently across the ribosome classes. Finally, we show that FPTs do not carry sequence hallmarks associated with protein-driven phase-separation propensity, suggesting that phase separation is not the likely functional context for the compositional biases we observe.

## Results

### Number of FPTs per ribosome varies substantially across domains

For each ribosome, we counted FPTs longer than 5 residues and longer than 20 residues. For mitoribosomes, we further filtered out portions responsible for targeting cytosole-translated proteins into mitochondria (see Methods). The number of protein tails is domain– and ribosome-specific (Fig. 1A–C). Mitoribosomes consistently showed the highest number of FPTs, followed by eukaryotic and bacterial ribosomes. While the differences in FPT sequence lengths on a median level are not very large across ribosome classes (Fig. 1D), mitoribosomes showed several FPTs significantly longer than any in eukaryotic and bacterial ribosomes.

**Figure 1:**
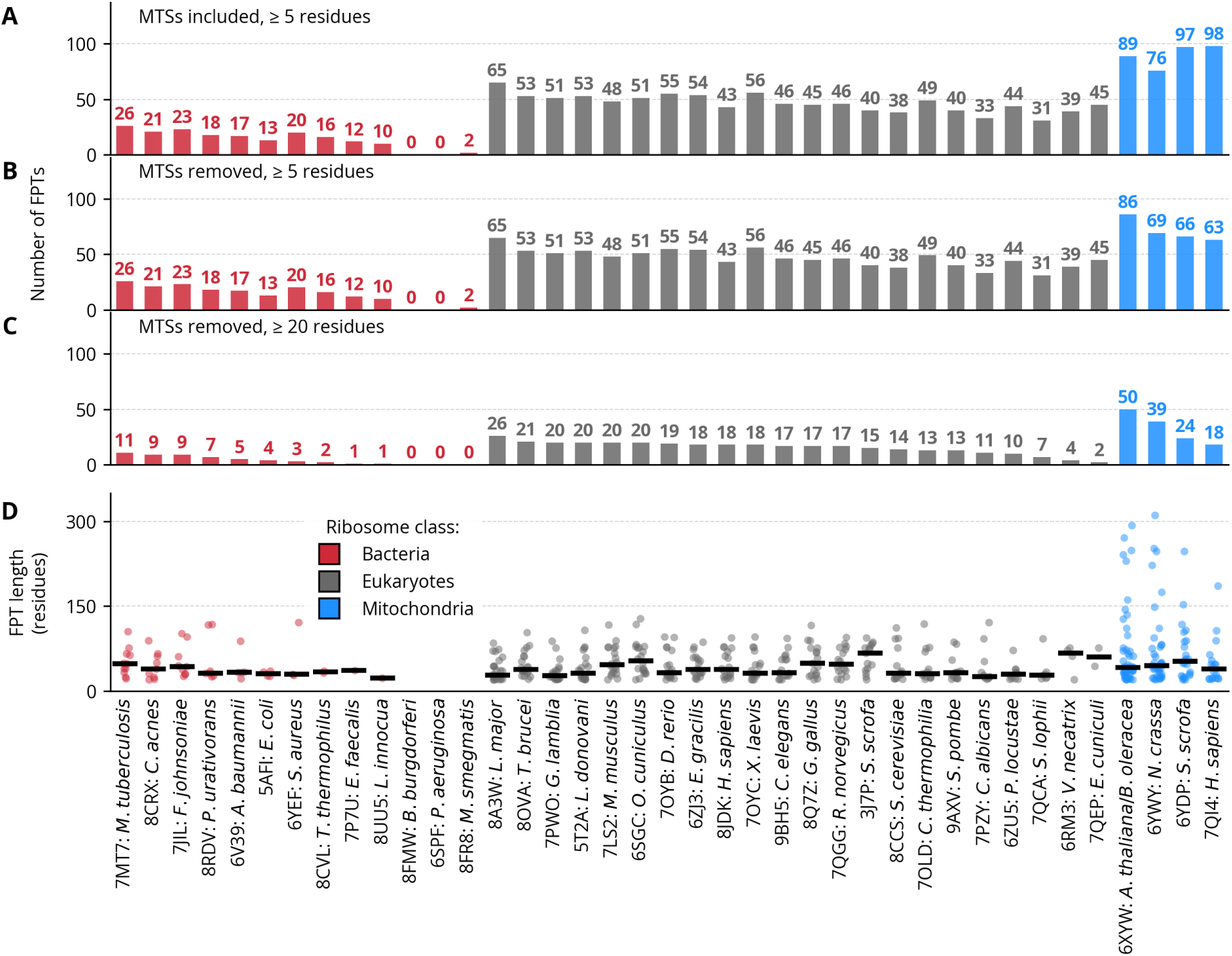
A) Number of flexible protein tails (FPTs) longer than 5 residues in each ribosome, mitochondrial targeting sequences (MTSs) included. B) Number of FPTs longer than 5 residues in each ribosome, MTSs removed. C) Number of FPTs longer than 20 residues in each ribosome. MTSs, removed. D) Individual dots show lengths of each FPT. The black line shows median FPT length. MTSs removed.

Eukaryotic and mitochondrial ribosomes contain many more ribosomal proteins than bacterial ribosomes overall (Fig. 2A) in the order bacteria *<* eukaryotes *<* mitochondria.

**Figure 2:**
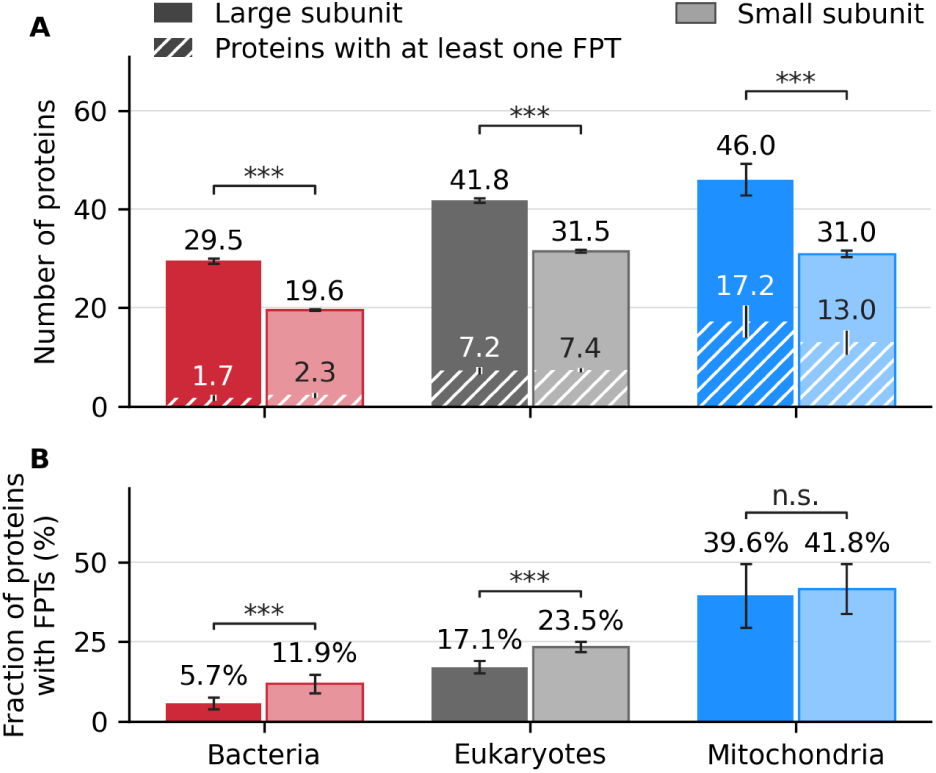
A) Average counts of ribosomal proteins and ribosomal proteins containing at least one FPT. B) Average percentage of proteins containing at least one FPT. Error bars denote SEM. Significance markers indicate p-values: ***: p *<* 0.001, n.s.: non-significant. Details of the statistical procedure are provided in the Methods section.

The percentage of FPT containing proteins is not constant across the ribosome classes, it also follows the same ordering with about 40% of mitoribosomal proteins containing FPTs. There is a trend toward higher FPT content in the small subunit across the ribosome classes, a difference of about 6 per-cent points for bacteria and eukaryotes (*p <* 0.001), and small, non-significant difference for mitochondria.

### FPTs are predominantly disordered

To assess the structural properties of FPTs, we predicted full-length protein structures using ESMFold^12^ and extracted per-residue pLDDT scores for the FPT regions. pLDDT serves as a proxy for disorder, expressing confidence in the predicted local three-dimensional structure; regions with pLDDT *<* 50 are considered disordered, while pLDDT *>* 70 corresponds to a generally correct backbone structure prediction.^13^

Per-residue pLDDT distributions across all ribosomes are shown in Fig. 3. The median pLDDT of FPTs for most ribosomes fell within the 50–70 range, indicative of moderate disorder of FPTs. Nonetheless, the distributions were broad and most ribosomes contained FPTs with a substantial fraction of residues with pLDDT *<* 50. This suggest a bifacial architecture of FPTs containing structured domains tethered with each other or to the ribosomes by unstructured linkers. The notable exceptions were several bacterial FPTs with median pLDDTs *>* 70 and *Staphylococcus aureus* approaching 100. Mitoribosomal FPTs showed the lowest pLDDT values overall, followed by eukaryotic and bacterial ribosomes, consistent with the trend observed for FPT length and number. *Giardia lamblia* was a single exception among eukaryotes, with a median pLDDT *<* 50 placing it closer to the mitoribosomal range.

**Figure 3:**
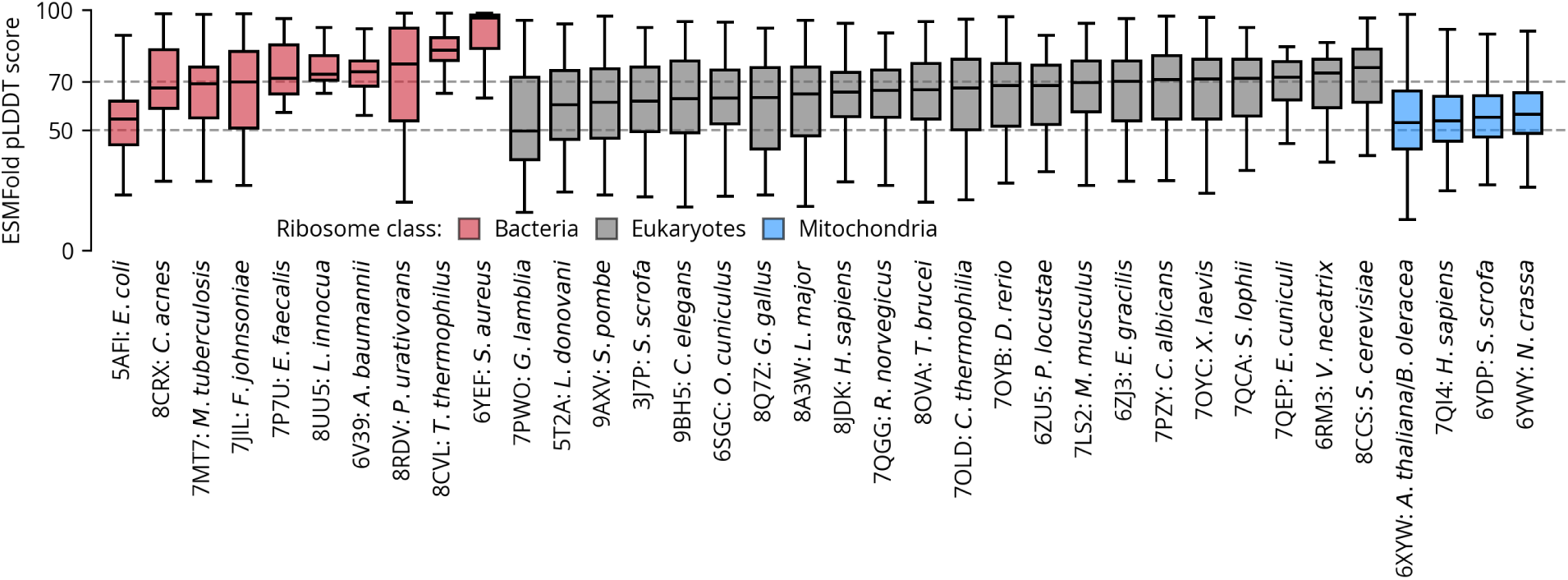
Per-residue pLDDT scores of the FPTs longer than 20 residues (MTSs removed) calculated using ESMFold. The boxplot box extends from the first quartile to the third quartile, with a line at the median. The whiskers extend from the box to the farthest data point lying within 1.5*×* the inter-quartile range from the box.

To further quantify disorder, secondary structure was assigned to ESMFold-predicted tertiary structures using DSSP, with the eight original secondary structure categories collapsed into three: coil, helix and strand (Fig. 4; see Methods). The fractional coil content of FPTs serves as an additional measure of disorder. With the exception of a subset of bacterial and a handful of eukaryotic ribosomes, the majority of FPTs across all ribosome types were assigned to the coil category, corroborating the pLDDT-based assessment of disorder.

**Figure 4:**
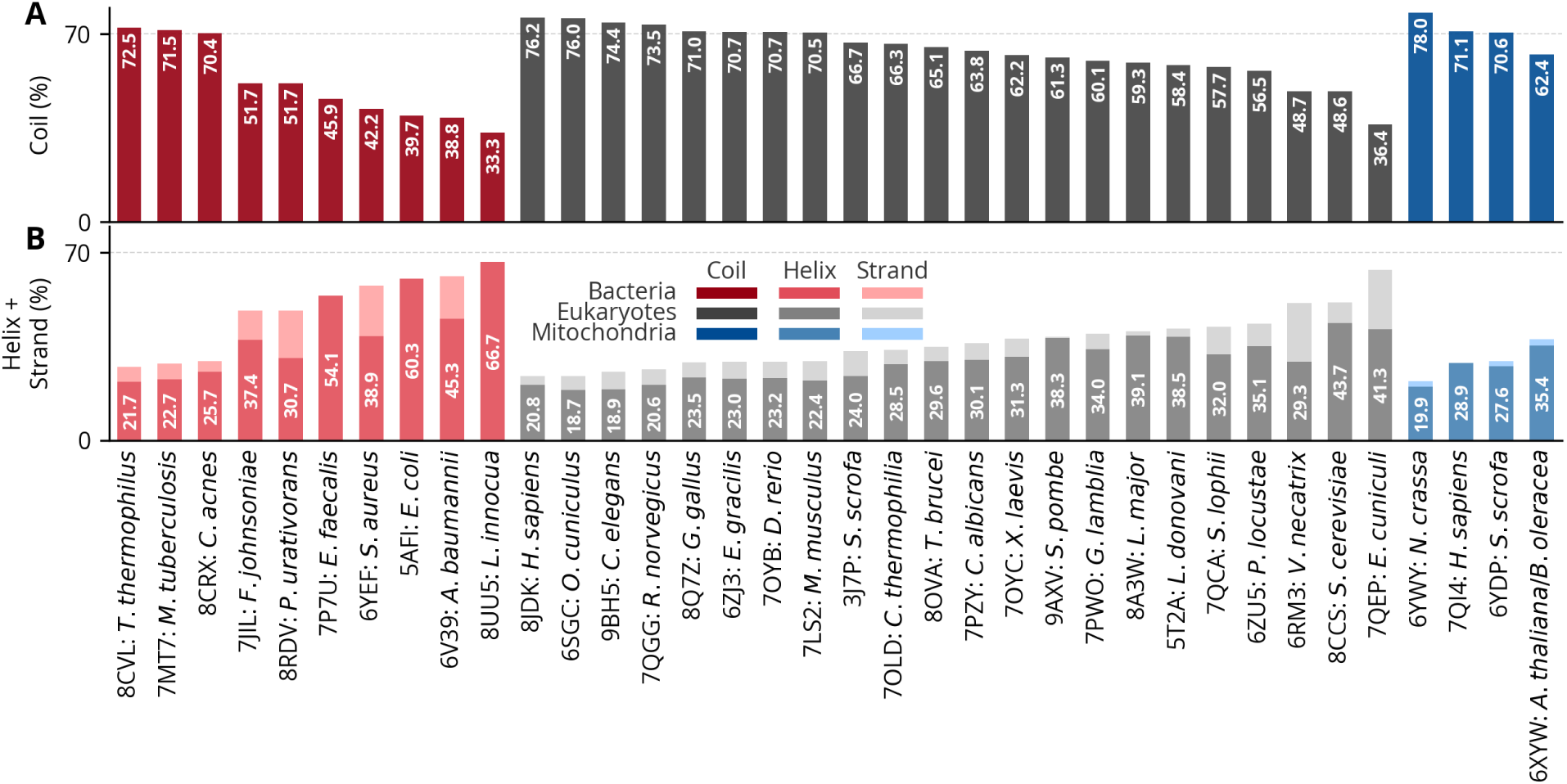
DSSP secondary structure content of FPTs aggregated per ribosome. The eight original secondary structure types were collapsed into three categories: coil, helix and strand.

As an orthogonal disorder evaluation we also used Metapredict (Fig. 5). As opposed to pLDDT, the higher the Metapredict score, the higher the disorder, with a sequence being considered disordered when the score is above 0.5. The results broadly agree with the pLDDT analysis, showing high disorder for mitochondria, most eukaryotes and some of the bacteria, there was however a disagreement in the case of *G. lamblia*, showing a median Metapredict score below 0.5, leaning towards a more ordered sequence, whereas the pLDDT analysis shows it as more disordered. The score distribution across its 20 FPTs is however quite wide.

**Figure 5:**
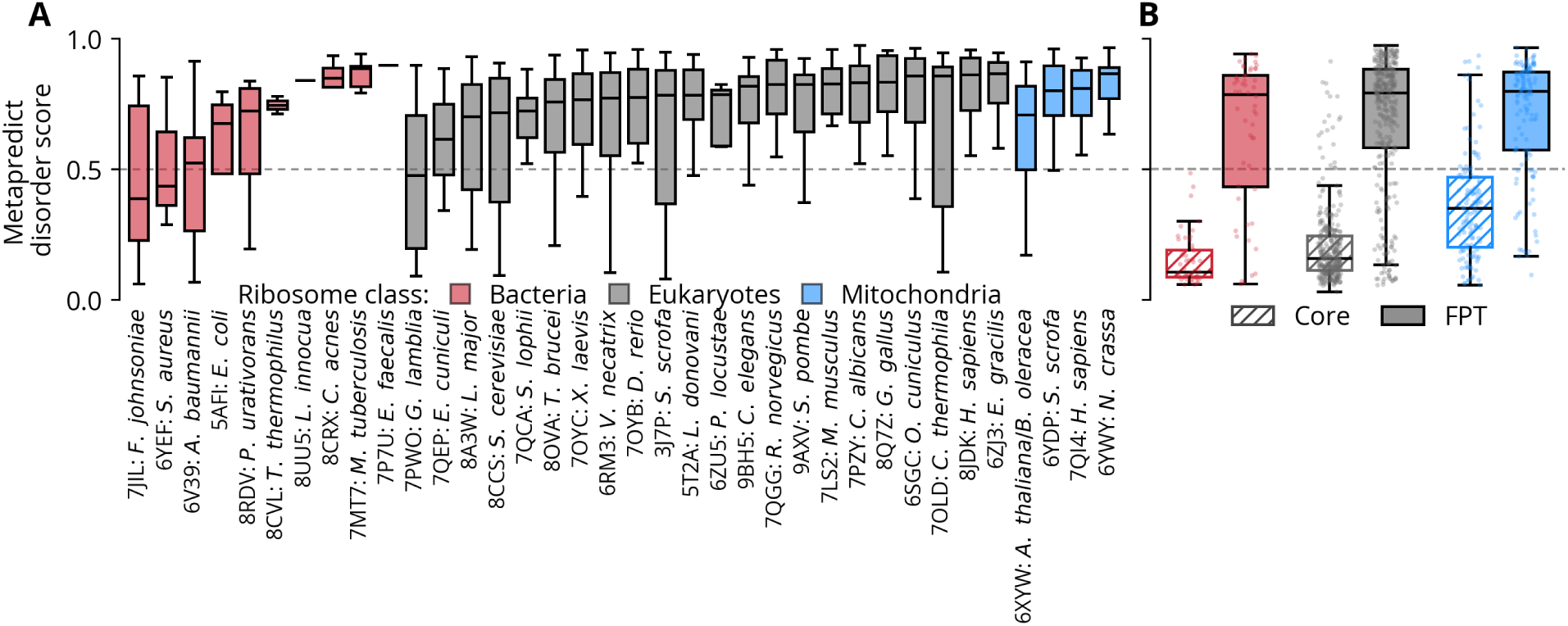
A) Boxplot of the Metapredict disorder score of FPTs by ribosome. B) Boxplot of the Metapredict disorder score of FPTs versus core regions of the ribosomes (non-FPT parts) by ribosome class. The individual points represent individual ribosomal proteins or FPTs. In both panels the boxplot box extends from the first quartile to the third quartile, with a line at the median. The whiskers extend from the box to the farthest data point lying within 1.5*×* the inter-quartile range from the box. Anything above the dashed line at 0.5 is considered disordered.

Across all three domains of life, FPTs showed consistently higher disorder than the corresponding protein core, reinforcing our results that ribosomal FPTs are genuinely disordered rather than merely poorly resolved regions.

### Amino acid composition of FPTs

We compared the amino acid composition of the FPTs of each ribosome class with the overall UniProtKB/Swiss-Prot background obtained from ExPASy (Fig. 6). Several features were shared by all three ribosome classes. Isoleucine was significantly depleted in bacterial, eukaryotic and mitochondrial FPTs, and aromatic residues were likewise depleted in all three classes (*p <* 0.001). Leucine was depleted and alanine enriched in bacterial and eukaryotic FPTs, with large and highly significant effects (*p <* 0.001), whereas mitochondrial FPTs showed only a small, non-significant trend in the same direction.

**Figure 6:**
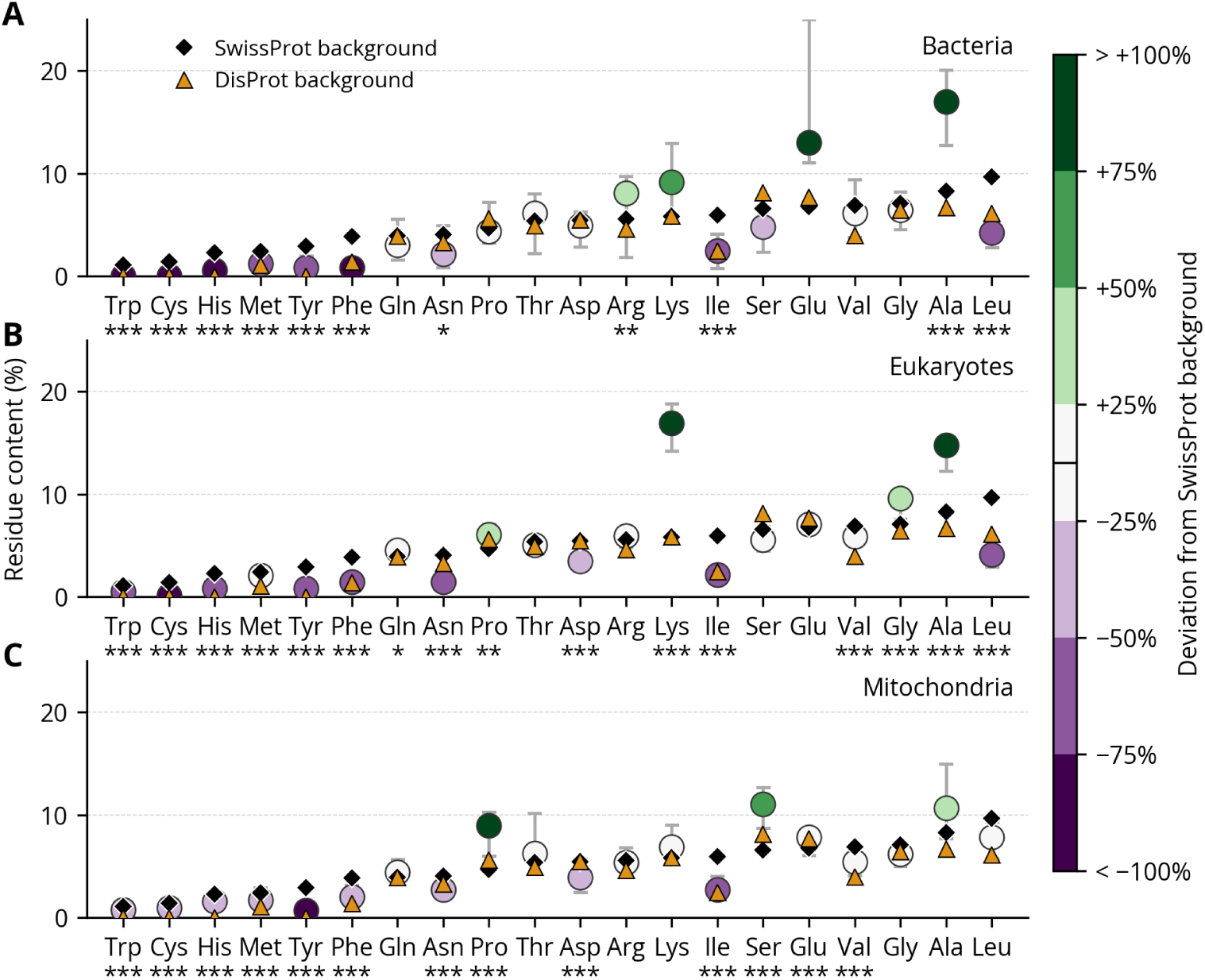
Amino acid composition of flexible protein tails (FPTs) longer than 20 residues (MTSs excluded). The black diamond denotes the Swiss-Prot proteome-wide amino acid frequency covering ordered and disordered proteins, the orange triangle denotes the amino acid frequency across DisProt representing IDPs and IDRs. Circles represent the median amino acid composition across FPTs of ribosomes of a given domain and are coloured according to enrichment (green) or depletion (purple) relative to the Swiss-Prot background. Bars represent 95% bootstrap confidence intervals. Significance markers below residue names indicate p-values relative to the Swiss-Prot background after Benjamini–Hochberg correction: ***: p *<* 0.001, **: p *<* 0.01, *: p *<* 0.05. Details of the statistical procedure are provided in the Methods section.

The ribosome classes differed in their charged and polar residue content. Bacterial FPTs showed a slight enrichment in arginine (*p <* 0.01) and a non-significant enrichment in lysine. In eukaryotic FPTs, arginine was unchanged relative to the background, whereas lysine was strongly and significantly enriched (median *∼*17%, *p <* 0.001). Mitochondrial FPTs showed no enrichment in arginine or lysine, but were significantly enriched in serine and proline (*p <* 0.001).

### FPTs are depleted in aromatic residues across all ribosome classes

To further characterise the sequence properties of the FPTs, we examined their median aromatic residue content (Tyr, Phe, Trp, and His combined) in relation to their median arginine content across all ribosomes in the dataset (Fig. 7). Dashed lines indicate the corresponding UniProtKB/Swiss-Prot proteome-wide background frequencies. No ribosome in the dataset showed enrichment of FPTs in aromatic residues relative to the background, while a handful of bacterial and eukaryotic ribosomes were enriched in arginine, consistent with the compositional analysis described above. Mitochondrial ribosomes likewise showed low aromatic content in FPTs and, unlike several bacterial and eukaryotic ribosomes, no enrichment in arginine. The uniformly low aromatic content of the unresolved terminal segments across all ribosome classes is noteworthy given the established role of aromatic residues in driving phase separation of disordered proteins.

**Figure 7:**
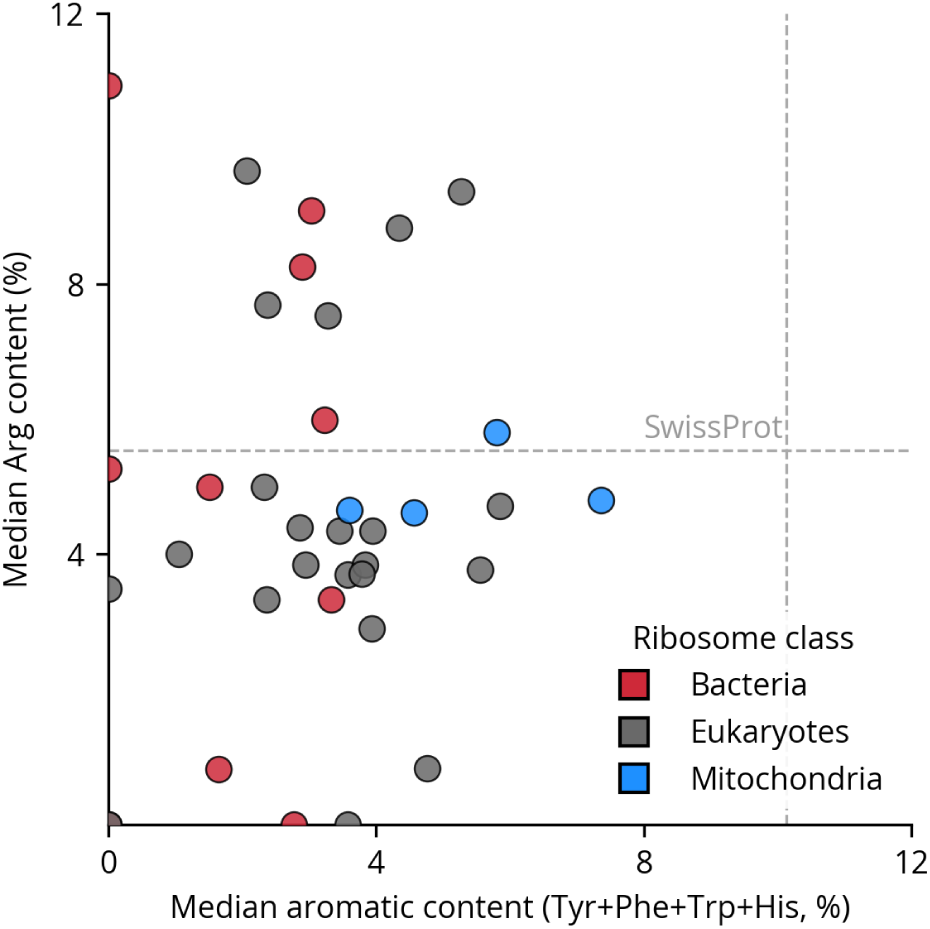
Aromatic residue content versus arginine content of ribosomal FPTs. Each point represents one ribosome, coloured by domain. Median Tyr, Phe, Trp and His content combined is shown on the x-axis and median Arg content on the y-axis. Dashed lines indicate Swiss-Prot proteome-wide background frequencies for aromatic residues (vertical) and arginine (horizontal).

### FPTs show low predicted phase-separation propensity

The predicted saturation concentrations of the FPTs are high and fall in the same range as those of the proteins that do not phase separate (Fig. 8). The median *c_sat_* of the FPTs from bacteria, eukaryotes and mitochondria falls in the 80 to 90 mg/mL range, comparable to the medians of the structured (NP) and disordered (ND) negative sets, and their interquartile ranges largely overlap with those of the negatives. In contrast, the condensation-prone proteins show markedly lower *c_sat_* values, with medians of roughly 55 mg/mL for CD and 70 mg/mL for DE, and a much broader distribution extending down to near zero. The bulk of the FPTs lies close to the highest *c_sat_* of the predictor’s training set (dashed line), the regime the model assigns to sequences that do not phase separate. Overall, the predictions do not suggest a propensity for protein-driven phase separation in any of the three FPT groups.

**Figure 8:**
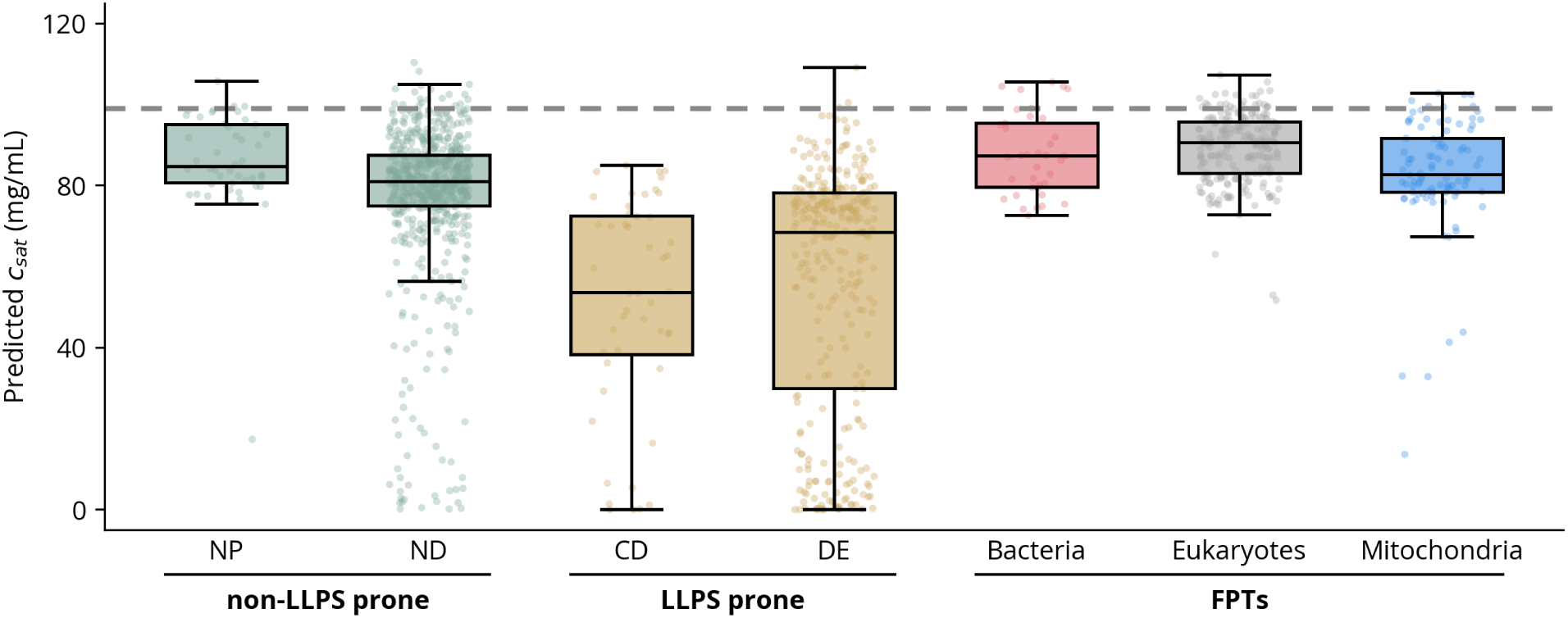
Predicted saturation concentration for FPTs longer than 20 residues (MTSs removed), liquid–liquid phase separation (LLPS) prone and non-LLPS prone proteins using tool by von Bülow *et al.*^14^ NP: non-LLPS prone structured proteins from the PDB. ND: non-LLPS prone DisProt proteins. DE: LLPS drivers exclusively. CD: proteins annotated as LLPS clients as well as drivers. Metapredict was used to select the longest IDR per protein for proteins from the comparison datasets (see Methods for details). The gray dashed line denotes the highest *c_sat_* of the proteins in the prediction tool’s training dataset. The boxplot box extends from the first quartile to the third quartile, with a line at the median. The whiskers extend from the box to the farthest data point lying within 1.5*×* the inter-quartile range from the box. The individual points represent individual IDRs or FPTs.

### Arg-Gly enrichment and mitochondria-specific Ala clustering in FPTs

Most amino acid group patterns in the FPTs were close to their shuffled expectation, with fold enrichments between 0.75*×* and 1.25*×* in all three ribosome classes (Fig. 9). The clearest signal was for the RGG motif, which was enriched in all three classes. RGG showed the strongest enrichment in eukaryotes (2.59*×*, ***), with similar or lower values in bacteria (2.46*×*) and mitochondria (1.90*×*), though neither of these was significant. RG/GR was also enriched in eukaryotes (1.65*×*, ***), but not in bacteria or mitochondria. KG/GK showed a weaker enrichment in eukaryotes (1.31x, ***), which suggests that the effect is stronger for arginine than for lysine. Among the group-level patterns, eukaryotic FPTs were enriched for positively charged residues next to glycine (Pos/Gly, 1.46*×*, ***) and for pairs of negatively charged residues (Neg-Neg, 1.25*×*, ***), and were depleted for glycine next to hydrophobic residues (Gly/Hyd, 0.74*×*, ***). Although alanine was abundant in the FPTs of all three ribosome classes, Ala–Ala pairs were enriched beyond compositional expectation only in mitochondria (1.54x, ***). Mitochondria also show Gly–Gly enrichment (1.51*×*, ***).

**Figure 9:**
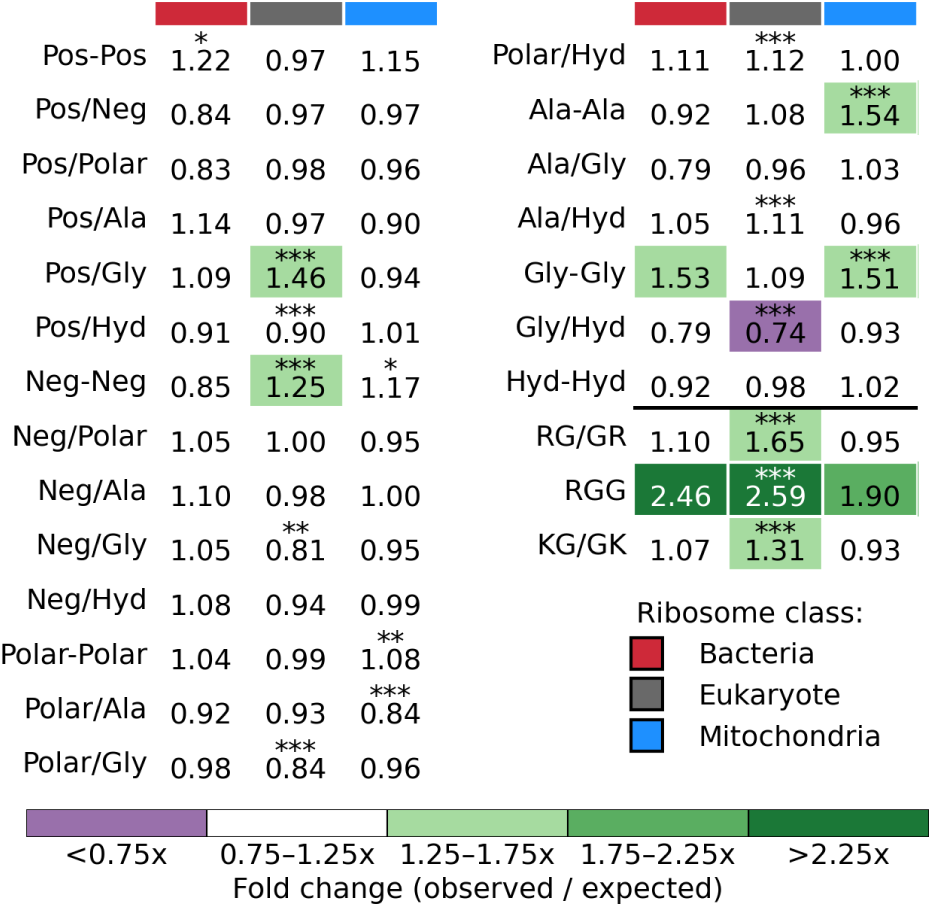
Enrichment of amino acid patterns in the FPTs of bacterial, eukaryotic and mitochondrial ribosomes. Cell values give the fold enrichment (observed / expected matches) of each pattern in each ribosome class. Expected counts were obtained from 20,000 shuffles of the amino acid positions within each FPT sequence, which preserves FPT composition. Cell colour indicates the fold-enrichment category. Patterns above the black line are combinations of amino acid groups (Pos, positively charged; Neg, negatively charged; Polar; Hyd, hydrophobic; Ala; Gly), where a hyphen denotes two residues from the same group and a slash denotes two residues from different groups in either order. Patterns below the line are the specific RG/GR, RGG and KG/GK motifs. Asterisks mark significance after Benjamini–Hochberg correction of empirical two-tailed p-values (*: p *<* 0.05, **: p *<* 0.01, ***: p *<* 0.001).

## Discussion

Despite forming a substantial protein content of up to thousands of residues on the ribosome surface, the flexible terminal regions of ribosomal proteins have received little attention, largely because their absence from experimental structures renders them effectively invisible to standard structural analyses. We characterised these flexible protein tails across ribosomes from bacteria, eukaryotes and mitochondria and found differences in their prevalence and length, class-specific sequence biases, and high levels of disorder. Below we first consider why the proportion of proteins carrying FPTs differs between ribosome classes, then what their composition implies about disorder and phase separation, and finally the basic residues of eukaryotic FPTs and the distinct signature of mitochondrial FPTs. Much of this discussion is necessarily hypothetical, and we indicate where this is the case.

The proportion of ribosomal proteins carrying FPTs follows the order bacteria *<* eukaryotes *<* mitochondria. One simple explanation is that this order tracks the protein content of the ribosome. Bacterial ribosomes are about one third protein and two thirds rRNA by mass,^15^ with a protein-to-rRNA ratio of about 1:2, whereas the large increase in r-protein mass in eukaryotes brings the ratio close to 1:1. ^16^ Mammalian mitoribosomes have inverted the bacterial ratio, at roughly 69% protein and 31% rRNA.^15^ Both eukaryotic and mitochondrial ribosomes gained protein mass partly through additional proteins and partly through N– and C-terminal extensions of conserved proteins,^17^ and such extensions are natural candidates for flexible, unresolved tails. Because bacterial ribosomal proteins interact mainly with rRNA, whereas eukaryote-specific segments engage in many protein–protein interactions,^16^ bacterial tails may also be more often constrained by the rRNA.

The same order is broadly the reverse of ribosome abundance and crowding, although the comparison is imperfect. Bacterial cells contain tens of thousands of ribosomes at high cytoplasmic densities during fast growth, ^18^ and bacterial polysomes show preferred, staggered ribosome orientations,^19^ although *in situ* data from a minimal bacterium suggest that only a minority of ribosomes are in closely assembled polysomes.^20^ Eukaryotic cytosolic ribosomes are on average less concentrated, though yeast cytoplasm approaches bacterial densities, and eukaryotic polysomes also form tightly packed helical or double-row assemblies.^21,22^ Mitoribosomes are far less numerous per cell and translate a tiny set of short transcripts, but they also form ordered polysomes on the inner membrane.^23,24^ Their local concentration in the matrix has not, to our knowledge, been measured. In bacterial and eukaryotic polysomes the small subunits face the polysome interior, close to the mRNA and to neighbouring ribosomes, whereas the large subunits face outward. ^19,21^ A simple steric-avoidance model would therefore predict fewer tails on the small subunit. Instead, the small subunit carries a slightly higher fraction of FPT-containing proteins in all three ribosome classes, so this line of thoughts is not supported and the inverse relation with crowding remains a correlation. Mapping which tails lie at inter-ribosome interfaces would test it directly, and other factors, such as eukaryote-specific assembly and regulatory functions or weaker purifying selection, could produce the same trend.

The compositional signature shared by all three classes is that of disordered regions. Isoleucine is strongly order-promoting and is depleted in FPTs of all classes, as expected for IDRs,^25^ and the pLDDT, DSSP and Metapredict analyses confirm that most FPTs are disordered or only marginally structured. Disorder was, however, more variable among bacterial ribosomes, some of which carry FPTs with lower predicted disorder than those of eukaryotic and mitochondrial ribosomes. Because bacterial ribosomes also have the fewest and shortest FPTs, per-ribosome estimates rest on fewer tails, and short tails may be more likely to retain partial secondary structure while still being unresolved. This heterogeneity is consistent with our operational definition of FPTs, which also includes flexibly tethered regions that are not fully disordered. It suggests that FPTs should not be assumed to behave as typical IDRs in bacteria. Alanine is considered a weakly disorder-promoting residue^25^ and is commonly found in disordered linkers because it lacks a bulky side chain that would drive hydrophobic collapse; it is significantly enriched in bacterial and eukaryotic FPTs, with only a non-significant trend in mitochondrial FPTs. Leucine depletion in bacterial and eukaryotic FPTs is likewise consistent with IDR character. These signals are therefore best interpreted as general compositional features of disordered regions rather than class-specific functional adaptations.

Neither the predicted saturation concentrations nor the sequence composition of FPTs suggest an intrinsic propensity for protein-driven phase separation. FPTs are strongly depleted in aromatic residues (Fig. 6, Fig. 7), more so than the DisProt background, which argues against the sticker-like interactions that drive self condensation of many disordered proteins.^26,27^ This also applies to RGG-containing regions: in FUS-family proteins, their arginines drive phase separation largely through interactions with tyrosines of partner prion-like domains,^28,29^ and FPTs lack such aromatic partners. Arginine-rich sequences without aromatic residues, such as the C9orf72 GR and PR dipeptide repeats, can nevertheless phase separate in vitro, ^30^ and RNA can promote phase separation of RG/RGG regions,^31^ so aromatic depletion argues against, but does not exclude, condensation of RGG-containing FPTs in the presence of RNA. The identity of the enriched basic residue points the same way: arginine-rich assemblies are more cohesive than lysine-rich ones, ^32,33^ and the strongly enriched basic residue in eukaryotic FPTs is lysine. This argument applies least to bacterial FPTs, which are modestly arginine-enriched, but these are also depleted in aromatic residues. The predictor models homotypic phase separation and cannot capture RNA-dependent heterotypic condensation, so our results argue against an intrinsic phase separation propensity of FPTs rather than excluding all roles in condensates, into which they might partition as clients.

The most striking class-specific signal is the large, significant lysine enrichment of eukaryotic FPTs (median *∼*17%), which is not accompanied by an arginine enrichment. The cell biology of eukaryotic ribosome assembly offers one candidate explanation. Eukaryotic r-proteins are synthesised in the cytoplasm, imported into the nucleus and assembled with rRNA in the nucleolus, and the resulting pre-ribosomes are exported again.^34,35^ Unassembled r-proteins are highly basic and aggregation-prone, and they depend on importins and dedicated chaperones that bind exposed basic segments.^36,37^ Their classical import signals are Lys/Arg-rich, are absent from bacterial homologues, and appear to have arisen by remodelling of rRNA-binding domains.^38^ Unlike mitochondrial presequences, these signals are not cleaved off and remain in the mature protein, but they appear to be largely buried in rRNA.^38,39^ They would therefore be expected to account for buried rather than surface-exposed lysines, and probably do not explain the enrichment in FPTs. A further, speculative function is charge compensation. Bacterial ribosomes bind hundreds of Mg^2+^ ions, and basic proteins alone do not neutralise the rRNA, ^40^ whereas free Mg^2+^ is estimated to be lower in eukaryotic (*∼*0.5–1 mM) than in bacterial (*∼*1.5–3 mM) cells.^41^ Lysine-rich surface tails might therefore partly substitute for Mg^2+^.

Despite the lack of arginine enrichment overall, RG/GR and RGG patterns occur more often than expected from composition in eukaryotic FPTs (Fig. 9). RGG showed comparable fold enrichments in bacterial, eukaryotic and mitochondrial FPTs (2.5*×*, 2.6*×* and 1.9*×*) but reached significance only in eukaryotes, whereas RG/GR and KG/GK were enriched only in eukaryotes. Because significance depends on sample size, these data do not show that RGG enrichment differs between ribosome classes. RG/RGG motifs are characteristic of disordered RNA-binding regions and are targets of arginine methylation; ^42,43^ a ribosomal example is the RG-repeat N-terminus of uS5 (rpS2), which is methylated by PRMT3.^44^ One possible division of labour is that lysine supplies diffuse positive charge, whereas arginine–glycine motifs provide more specific, flexible contacts with RNA, but this is speculative. The RGG enrichment may reflect transient, low-specificity contacts of the arginine side chains with exposed rRNA, including expansion segments in eukaryotes. Sequence composition alone, however, cannot distinguish selection for such contacts from a consequence of tail basicity or from earlier roles in nuclear import and assembly.^35^ The eukaryote-specific RG/GR enrichment may likewise reflect these demands or PTM-based regulation. We consider it more likely that FPTs engage RNA and partner proteins heterotypically than that they condense on their own.

FPTs could also serve as recognition sites for chaperones, but two groups of ribosome-related chaperones need to be distinguished. The first are dedicated chaperones of unassembled ribosomal proteins. In eukaryotes, several of these capture newly synthesised ribosomal proteins through eukaryote-specific N-terminal extensions or loops, protecting them from aggregation and escorting them to the assembly site,^35,45^ which is consistent with the greater prevalence of such extensions in eukaryotic ribosomes. Dedicated chaperones are known for only a small fraction of ribosomal proteins,^37^ so additional FPT-binding factors may remain to be identified. Because chaperone-binding segments are often later involved in rRNA binding,^45^ it is unclear whether the tails that remain unresolved in mature ribosomes overlap with these sites. The second group comprises ribosome-associated protein biogenesis factors, which bind mainly near the exit tunnel of the large subunit and assist the folding, modification and targeting of nascent chains.^46^ These factors act on the emerging nascent chain and not on the ribosomal proteins themselves, so they are not expected to recognise FPTs, and, since they bind the large subunit, they would not explain the slightly higher small-subunit fraction of FPT-containing proteins. For mitoribosomes, which have the highest proportion of FPT-bearing proteins, and for bacterial ribosomes, which have the lowest, we are not aware of comparable dedicated ribosomal-protein chaperones, and whether FPT content reflects chaperone-mediated processes in these systems is unknown. This possibility could be tested by comparing FPT-bearing proteins with known chaperone clients and with mapped chaperone-binding segments.

Mitochondrial FPTs differ from the other two classes. After correction for cleaved presequences they show no enrichment of arginine or lysine and, in line with the broader pattern of mostly null or small compositional differences, no significant deviation in leucine, so the basic-residue explanations above do not extend to them. They are, however, enriched in proline and serine, both strongly disorder-promoting residues, and Ala–Ala pairs occur more often than expected from composition in mitochondria only (1.5*×*; Fig. 9). Alanine content is a generic feature of disordered regions, but this composition cannot by itself explain clustering beyond what is expected from it. Gly–Gly pairs were also enriched, so clustering of small residues in general may be involved. We can offer three non-exclusive and speculative explanations. First, the high serine content may reflect phosphorylation sites: mitoribosomal proteins are phosphorylated at serine, threonine and tyrosine residues, ^47^ and phosphorylation of mitoribosomal components has been proposed to regulate mitochondrial translation,^48^ although threonine, the other main phosphoacceptor, is not enriched. Second, the combined enrichment of proline, serine and alanine resembles synthetic Pro/Ala/Ser polypeptides that form hydrophilic, expanded random coils,^49^ which would suit flexible, solvent-exposed tails. Third, alanine runs in disordered regions can form transient helices that mediate protein– protein interactions,^50^ which would suit mitoribosomal extensions that mainly contact other mitochondria-specific proteins.

Several limitations apply. FPTs are defined operationally by their absence from experimental models, so they include genuinely disordered sequences as well as flexibly tethered segments that retain some structure. The compositional analysis treats ribosomes as the unit of observation, whereas the pattern analysis pools FPTs, so the two differ in sensitivity, and differences in significance between classes, particularly for the mitochondrial ribosomes, which are few, should be read cautiously. Most of the explanations above are untested, but several yield concrete predictions: mapping which tails lie at inter-ribosome interfaces, comparing FPT-bearing proteins with known chaperone clients, measuring the distance of tails to exposed rRNA and expansion segments and simulating their contacts, mapping phosphorylation sites onto mitoribosomal FPTs, and testing the helical propensity of Ala–Ala-containing segments. Taken together, our results show that the unresolved tails of ribosomal proteins are neither compositionally random nor uniform across the domains of life, and they identify several specific hypotheses for their function.

## Methods

### Dataset

The starting point for the dataset was the collection of ribosome structures from the PDB compiled by W-lodarski.^51^ From these, only ribosome structures containing both subunits determined at a resolution of 4 Å or better were considered. For these entries, both the mm-CIF files (from which FASTA sequences of proteins resolved in the structure were extracted) and the full respective FASTA sequences themselves were taken from the Protein Data Bank. A small number of ribosomal protein sequences contained ambiguous residues (denoted X in the FASTA sequence) and were excluded from the analysis. Also excluded were any non-ribosomal proteins (associated enzymes etc.) The entries for *Archaea* and chloroplasts were removed from the dataset as only two and one remained respectively. A few further entries outside the W-lodarski set that fulfilled all criteria were added to the dataset. The final dataset contains 22 eukaryotic, 13 bacterial and 4 mitochondrial ribosomes, of which 3 bacterial ribosomes contained no FPTs longer than 20 residues and did not feature in any further FPT analyses. Table 1 contains the full list of ribosomes used.

**Table 1:** Summary of the ribosome structures selected for analysis.

| **PDB ID** | **Organism** | **Ribosome class** | **Resolution (Å)** |
|---|---|---|---|
| 6V39 | *A. baumannii* | Bacteria | 3.04 |
| 6YEF | *S. aureus* | Bacteria | 3.20 |
| 8CRX | *C. acnes* | Bacteria | 2.78 |
| 5AFI | *E. coli* | Bacteria | 2.90 |
| 7P7U | *E. faecalis* | Bacteria | 3.10 |
| 7JIL | *F. johnsoniae* | Bacteria | 2.80 |
| 8UU5 | *L. innocua* | Bacteria | 3.00 |
| 8FR8 | *M. smegmatis* | Bacteria | 2.76 |
| 7MT7 | *M. tuberculosis* | Bacteria | 2.71 |
| 8RDV | *P. urativorans* | Bacteria | 2.60 |
| 8CVL | *T. thermophilus* | Bacteria | 2.30 |
| 6SPF | *P. aeruginosa* | Bacteria | 3.34 |
| 8FMW | *B. burgdorferi* | Bacteria | 2.86 |
| 7PZY | *C. albicans* | Eukaryotes | 2.32 |
| 9BH5 | *C. elegans* | Eukaryotes | 2.63 |
| 7OLD | *C. thermophilia* | Eukaryotes | 3.00 |
| 7OYB | *D. rerio* | Eukaryotes | 2.40 |
| 7QEP | *E. cuniculi* | Eukaryotes | 2.70 |
| 7PWO | *G. lamblia* | Eukaryotes | 2.75 |
| 9AXV | *S. pombe* | Eukaryotes | 2.40 |
| 6ZJ3 | *E. gracilis* | Eukaryotes | 3.15 |
| 8Q7Z | *G. gallus* | Eukaryotes | 2.50 |
| 8JDK | *H. sapiens* | Eukaryotes | 2.26 |
| 5T2A | *L. donovani* | Eukaryotes | 2.90 |
| 8A3W | *L. major* | Eukaryotes | 2.89 |
| 7LS2 | *M. musculus* | Eukaryotes | 3.10 |
| 6SGC | *O. cuniculus* | Eukaryotes | 2.80 |
| 6ZU5 | *P. locustae* | Eukaryotes | 2.90 |
| 7QGG | *R. norvegicus* | Eukaryotes | 2.86 |
| 8CCS | *S. cerevisiae* | Eukaryotes | 1.97 |
| 7QCA | *S. lophii* | Eukaryotes | 2.79 |
| 3J7P | *S. scrofa* | Eukaryotes | 3.50 |
| 8OVA | *T. brucei* | Eukaryotes | 2.47 |
| 6RM3 | *V. necatrix* | Eukaryotes | 3.40 |
| 7OYC | *X. laevis* | Eukaryotes | 2.40 |
| 6XYW | *A. thaliana/B. oleracea* | Mitochondria | 3.86 |
| 7QI4 | *H. sapiens* | Mitochondria | 2.20 |
| 6YWY | *N. crassa* | Mitochondria | 3.05 |
| 6YDP | *S. scrofa* | Mitochondria | 3.00 |

To identify FPTs absent from the experimental structure, we compared the full sequences of the ribosomal proteins to the sequences of the parts that are resolved in the experimental structures. Internal unresolved regions were not considered. Segments shorter than 20 residues were excluded from the analyses, as very short unresolved termini likely reflect local thermal disorder at protein termini rather than extended flexible regions of potential functional significance. While this threshold is somewhat arbitrary, the amino acid compositional signals, secondary structure content and pLDDT reported here were robust to varying this cutoff between 15 and 25 residues, with no qualitative differences that would affect the conclusions.

Mitoribosomal FPTs were further processed to exclude mitochondrial targeting sequences. Because MTSs are cleaved during protein maturation, any portion of an FPT extending into an MTS represents a region absent from mature mitoribosomal proteins rather than an unresolved segment. To correct for this, we used TargetP 2.0^52^ to predict MTS boundaries for each protein and removed any predicted MTS residues overlapping with the FPT region. Of the 360 mitoribosomal FPTs evaluated, 169 (46.9%) were unaffected by this step, 131 (36.4%) were trimmed to remove overlapping MTS residues, and 60 (16.7%) were excluded entirely as their full length fell within the predicted MTS. While MTS predictions from an alternative tool MitoFates^53^ broadly agreed with TargetP 2.0, minor discrepancies arose in a small subset of cases. Comparisons against available experimental data consistently supported the accuracy of the TargetP 2.0 predictions.

Annotations of subunit affiliation were inferred from Ban *et al.* ribosomal protein names^54^ found through RiboXYZ^55^ where available, otherwise through regular expression matching of the proteins names from the PDB and a small number of cases was annotated manually.

### Subunit differences

Differences between the large and small ribosomal subunits were tested separately for each ribosome class, for two metrics: the number of ribosomal proteins per PDB entry and the percentage of those proteins with at least one FPT. Because the two subunits of a given entry belong to the same ribosome, they were treated as paired observations. For each entry, the difference between the large and small subunit was computed, and the mean difference was tested against zero using a paired bootstrap: entries were resampled with replacement 10,000 times, and the mean difference was recomputed for each resample. The two-sided p-value was defined as twice the smaller of the proportions of bootstrap means at or below zero and at or above zero, p-values were corrected for multiple comparisons using the Benjamini–Hochberg false discovery rate procedure.^56^

### Amino acid composition

Amino acid composition was calculated for each terminal missing segment by computing the fractional occurrence of each of the 20 standard amino acids relative to segment length. For each ribosome, per-residue fractions were averaged across all qualifying terminal segments to produce a single per-ribosome mean for each amino acid, treating the ribosome as the unit of observation. Compositions were compared to proteome-wide background frequencies derived from the UniProtKB/Swiss-Prot database^57^ with per-amino-acid frequencies obtained from the ExPASy proteomics server. ^58^ The compositions were also compared to the median amino acid composition calculated from the complete DisProt dataset, 2026 06 release.^59^

Statistical significance of compositional deviations from the Swiss-Prot background was assessed using a shifted bootstrap procedure. For each amino acid and each ribosome class, the observed median content was computed and a bootstrap distribution of the median was generated by resampling ribosomes with replacement (10,000 iterations). To construct a null distribution centred on the Swiss-Prot background frequency rather than the observed median, each bootstrap distribution was shifted by subtracting the observed median and adding the Swiss-Prot value. Two-tailed p-values were computed as twice the proportion of the shifted null distribution falling at least as far from the Swiss-Prot value as the observed median. Bootstrap confidence intervals were computed separately as the 2.5th and 97.5th percentiles of the unshifted bootstrap distribution, providing an estimate of uncertainty around the observed median independent of the hypothesis test. To account for the simultaneous testing of all 20 amino acids within each group, p-values were corrected for multiple comparisons using the Benjamini–Hochberg false discovery rate procedure. ^56^ Note that significance markers reflect a bootstrap shift test on the median; apparent overlap between a confidence interval and the Swiss-Prot background value (dashed line) can co-occur with a significant result when the bootstrap distribution is asymmetric.

To assess the propensity of terminal missing segments for liquid-liquid phase separation, mean aromatic residue content (Tyr, Phe, Trp and His combined) and mean arginine content were calculated per ribosome by averaging across all qualifying terminal segments and plotted against each other. Reference lines representing the Swiss-Prot background are drawn at 10.13% for aromatic content and 5.53% for arginine content.

### pLDDT

ESMFold predicted structures were generated for all ribosomal protein chains containing FPTs longer than 20 residues using the ESMFold v1 model^12^ accessed via the Hugging Face transformers library. Predicted Local Distance Difference Test (pLDDT) scores were extracted from the model output and scaled to the 0–100 range. For each chain, the pLDDT scores were averaged across valid atoms using the atom37 existence mask to obtain a single confidence value per residue. pLDDT scores reflect the model’s confidence in the predicted local structure, with low scores (below 50) indicating regions lacking a stable tertiary structure and serving as a proxy for intrinsic disorder. Analysis was restricted to residues belonging to FPTs.

### Secondary structure content

Secondary structure was assigned to the terminal disordered segments using a two-step approach. For each ribosomal protein with unresolved terminal residues, the complete protein sequence was submitted to ESMFold to obtain a full-length predicted structure. Secondary structure was then assigned to all residues of each predicted structure using PyDSSP, a Python implementation of the DSSP algorithm.^60^ The eight-state DSSP assignments were collapsed into three categories for summary statistics: helix (*α*-helix, 3_10_-helix, and *π*-helix), strand (*β*-strand and *β*-bridge), and coil (loop/irregular and all remaining states). Secondary structure compositions were computed per ribosome by aggregating residue-level assignments across all qualifying FPTs from all chains, and the fraction of coil residues was used as a proxy for intrinsic disorder.

### Metapredict

As additional evidence of the disordered character of ribosomal FPTs, we used Metapredict,^61^ a data-driven predictor directly based on amino acid sequence, bypassing the structure prediction process. Thus Metapredict provides an orthogonal assessment with ESMFold which we use to infer disorder from confidence metrics.

For each ribosomal protein with an FPT, we computed the mean Metapredict disorder score separately over the FPT and over the ribosome core and we show these results aggregated per ribosome class. We also show FPT Metapredict scores for each ribosome separately.

### Phase separation propensities by PSpred

We predicted saturation concentration of the FPTs as a measure of phase-separation propensity using the prediction tool developed by von Bülow *et al.*^14^ To place these predictions in context, we compared them with sequences from the datasets by Pintado-Grima *et al.*^62^ As negative examples, with no evidence of phase separation, we used the NP set (structured proteins from the PDB) and the ND set (disordered proteins from DisProt). As positive examples, we used the DE set (proteins annotated exclusively as LLPS drivers) and the CD set (proteins annotated as both clients and drivers). NP and ND were pooled as the negative reference, and DE and CD as the positive reference. For each protein we used only its longest intrinsically disordered region, identified with Metapredict. Because the predictor was trained on sequences of at least this length, we only retained IDRs longer than 30 residues. The predictor was then applied to these IDRs, and the resulting predictions were compared with those obtained for the FPTs.

### Sequence patterns

To evaluate the statistical enrichment of specific amino acid patterns and motifs within the FPTs, we implemented a permutation– and bootstrap-based computational pipeline. For each ribosome class, the observed frequency of each pattern or motif was quantified across all valid FPT sequences. To establish a null expectation, background distributions were generated by randomly shuffling the amino acid positions of each FPT sequence 20,000 times while preserving individual sequence composition. Fold enrichment was calculated as the ratio of observed to expected matches. Empirical two-tailed *p*-values were derived by comparing the total observed match count against the null distribution generated from the shuffled sequences. Raw *p*-values were adjusted using the Benjamini-Hochberg procedure^56^ to control the false discovery rate across multiple comparisons.

## Acknowledgement

We acknowledge VSB – Technical University of Ostrava, IT4Innovations National Supercomputing Center, Czech Republic, for awarding this project access to the LUMI supercomputer, owned by the EuroHPC Joint Undertaking, hosted by CSC (Finland) and the LUMI consortium through the Ministry of Education, Youth and Sports of the Czech Republic through the e-INFRA CZ (grant ID: 90254). HM acknowledges the grants of Specific university research (A1 FCHI 2026 001 and A2 FCHI 2026 042). XG acknowledges funding from an EMBO Postdoctoral Fellowship (ALTF 1472-2025).

## Authors’ contribution

HM: data curation, formal analysis, visualisation, results interpretation, writing – original draft, funding acquisition; RK: data curation; XG: formal analysis, results interpretation, funding acquisition; MK: conceptualization, supervision, results interpretation, funding acquisition. All authors reviewed and edited the manuscript.

## Data and software availability

The dataset of FPTs and python scripts necessary to generate all figures are available on https://github.com/kolarlab/mcgrath-fpts.

